# Metabolic control of adaptive β-cell proliferation by the protein deacetylase SIRT2

**DOI:** 10.1101/2024.02.24.581864

**Authors:** Matthew Wortham, Bastian Ramms, Chun Zeng, Jacqueline R. Benthuysen, Somesh Sai, Dennis P. Pollow, Fenfen Liu, Michael Schlichting, Austin R. Harrington, Bradley Liu, Thazha P. Prakash, Elaine C Pirie, Han Zhu, Siyouneh Baghdasarian, Johan Auwerx, Orian S. Shirihai, Maike Sander

## Abstract

Selective and controlled expansion of endogenous β-cells has been pursued as a potential therapy for diabetes. Ideally, such therapies would preserve feedback control of β-cell proliferation to avoid excessive β-cell expansion and an increased risk of hypoglycemia. Here, we identified a regulator of β-cell proliferation whose inactivation results in controlled β-cell expansion: the protein deacetylase Sirtuin 2 (SIRT2). *Sirt2* deletion in β-cells of mice increased β-cell proliferation during hyperglycemia with little effect in homeostatic conditions, indicating preservation of feedback control of β-cell mass. SIRT2 restrains proliferation of human islet β-cells cultured in glucose concentrations above the glycemic set point, demonstrating conserved SIRT2 function. Analysis of acetylated proteins in islets treated with a SIRT2 inhibitor revealed that SIRT2 deacetylates enzymes involved in oxidative phosphorylation, dampening the adaptive increase in oxygen consumption during hyperglycemia. At the transcriptomic level, *Sirt2* inactivation has context-dependent effects on β-cells, with *Sirt2* controlling how β-cells interpret hyperglycemia as a stress. Finally, we provide proof-of-principle that systemic administration of a GLP1-coupled *Sirt2*-targeting antisense oligonucleotide achieves β-cell selective *Sirt2* inactivation and stimulates β-cell proliferation under hyperglycemic conditions. Overall, these studies identify a therapeutic strategy for increasing β-cell mass in diabetes without circumventing feedback control of β-cell proliferation.

## INTRODUCTION

Insufficient mass of functional β-cells underlies both type 1 and type 2 diabetes. Type 1 diabetes (T1D) results from autoimmune destruction of the majority of β-cells, with residual β-cells retaining some degree of function^1^. Type 2 diabetes (T2D) predominantly involves dysfunction of β-cells that have been challenged to sustain a high level of compensatory insulin secretion due to insulin resistance^2,3^. As the reduction of β-cell mass is the principal cause of hyperglycemia in T1D, and β-cell dysfunction in T2D is largely reversible by normalization of β-cell workload^4^, therapeutically increasing β-cell mass through proliferation has been proposed for treating both major types of diabetes.

Studies investigating therapeutic targets for expanding β-cell mass have identified several compounds that stimulate proliferation of adult human β-cells^5–8^. Recent findings from several groups converged upon mechanisms involving inhibition of the kinase DYRK1A, with further increases of β-cell proliferation achieved using combination treatment with DYRK1A inhibitors and other agents including GLP1^6^ or inhibitors of the TGFβ pathway^5^. The effectiveness of β-cell expansion therapies in preclinical models generally supports the feasibility of this approach^6,9^, which is currently awaiting safety and efficacy results from clinical trials (NCT05526430)^10^.

Given the advances in achieving effective β-cell expansion via experimental compounds, refinement of such approaches will address remaining concerns including mitogenic effects in non-β cells^11^ and the possibility of insulinoma formation or hypoglycemia. Feedback control of cell expansion is a desirable feature of any cellular regeneration therapy. One mechanism controlling precise expansion of β-cell mass in rodents is the glucose dependence of adaptive β-cell proliferation^12^, which couples β-cell proliferation to increased β-cell workload indicative of a shortfall of insulin. While several experimental compounds have demonstrated effectiveness in stimulating human β-cell proliferation^5,6^, the proposed therapies circumvent feedback regulation normally responsible for indicating that a set point for β-cell mass has been achieved. Preserving feedback regulation of proliferation during therapeutic β-cell expansion would have the advantage of generating the precise number of β-cells required for glycemic control.

Here we have identified a conserved mechanism that restrains β-cell proliferation in a glucose-dependent manner: metabolic control of proliferation via the protein deacetylase Sirtuin2 (SIRT2). SIRT2 is a member of the Sirtuin family of deacetylases that use nicotinamide adenine dinucleotide (NAD^+^) as a cosubstrate^13^. Through genetic, pharmacological, and proteomic approaches, we show that SIRT2 is a suppressor of β-cell proliferation at elevated glucose concentrations. SIRT2 inactivation enhances β-cell proliferation in vivo and ex vivo only at glucose concentrations above the normal glycemic set point, thereby preserving feedback control of β-cell expansion. Preclinical studies employing β-cell-specific *Sirt2* knockdown in vivo recapitulate this effect, establishing proof-of-principle for selectively and safely increasing β-cell numbers.

## RESULTS

### SIRT2 restrains proliferation of mouse and human β-cells ex vivo

β-cell proliferation in the postnatal period underlies developmental expansion of β-cells in both rodents and humans. This process generates sufficient β-cell mass for controlling glycemia in adulthood, at which point β-cell proliferation declines precipitously^14,15^. Following developmental maturation, rodent but not human β-cells maintain the capacity for adaptive proliferation in response to perceived insulin deficits. While this adaptive response is robust in young (1-2-month-old) animals, adaptive β-cell proliferation is progressively attenuated with age^16–18^. Given the age dependence of both basal and adaptive β-cell proliferation, we reasoned that mechanisms controlling proliferation could be identified through assessment of age-regulated features of the β-cell. To this end, we performed quantitative in vivo proteomics of islets from 1-month-old and 1-year-old mice^19^ corresponding to ages with starkly different capacities for adaptive β-cell proliferation^16,18^ and rates of basal β-cell proliferation^20^. Of differentially abundant proteins, we selected those with established roles in proliferation for follow-up studies, initially focusing on SIRT2 based on its known role in suppressing proliferation of immune cells and of various transformed cell types^21,22^. SIRT2 protein was elevated in islets from 1-year-old compared to 1-month-old mice (Supplementary Figure 1A), indicating an inverse association between SIRT2 abundance and capacity for β-cell proliferation. Immunofluorescence staining identified SIRT2 in both α-cells and β-cells in mouse pancreas and predominantly in β-cells in human islets (Figure 1A). The presence of SIRT2 in mouse and human β-cells as well as its age-dependent increase in abundance led us to predict that SIRT2 is a conserved regulator of β-cell proliferation.

**Figure 1.**
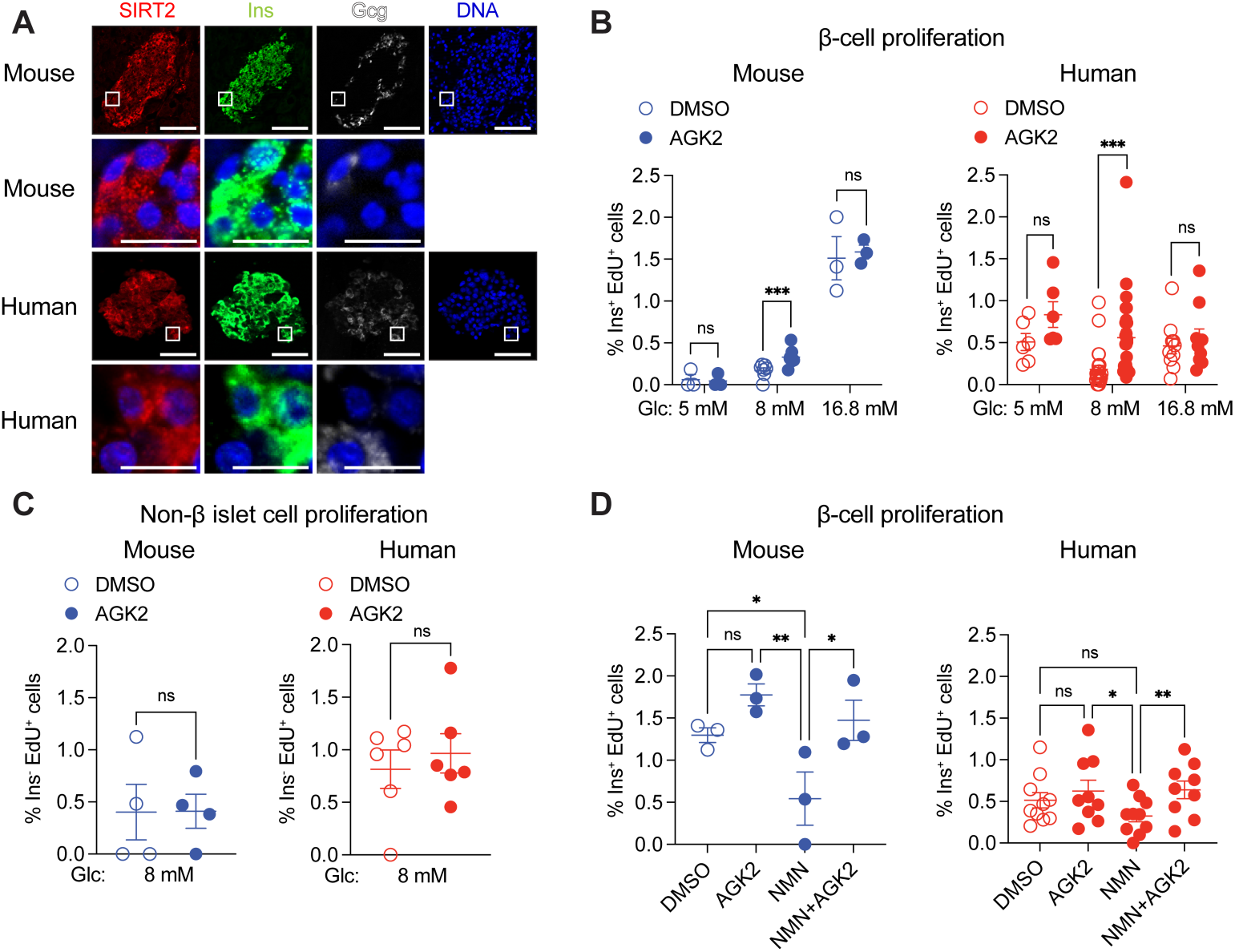
SIRT2 inhibition stimulates β-cell proliferation in a glucose-dependent manner in cultured mouse and human islets. (**A**) Immunofluorescence staining for the indicated proteins in representative mouse pancreata and human islet sections. Enlarged images for indicated areas including DNA overlay are shown below each channel. Scale bars, 50 μm (top) and 10 μm (enlarged images at bottom). Sections were counterstained for DNA using DAPI. (**B**) Quantification of β-cell proliferation as percentage of insulin-expressing cells positive for EdU in isolated mouse (blue, *n* = 3-7 islet preparations/group) and human (red, *n* = 6-25 islet preparations/group) islets after DMSO or AGK2 treatment during culture with the indicated glucose concentrations. (**C**) Quantification of non-β islet cell proliferation as percentage of insulin-negative cells positive for EdU in isolated mouse (blue, *n* = 4 islet preparations/group) and human (red, *n* = 6 islet preparations/group) islets after DMSO or AGK2 treatment. (**D**) Quantification of β-cell proliferation as percentage of insulin-expressing cells positive for EdU in isolated mouse (blue, *n* = 3 islet preparations/group) and human (red, *n* = 9-10 islet preparations/group) islets after treatment with DMSO, AGK2, NMN (an NAD^+^ precursor) or combinations thereof at 16.8 mM glucose. Data are shown as mean ± SEM. Statistical differences were calculated using a multiple paired t-tests (B) or paired t-test (C) to determine statistical differences between two groups. A one-way ANOVA with Tukey post hoc analysis was performed to analyze statistical differences between three or more groups (D). \**p*<0.05, \*\**p*<0.01, \*\*\**p*<0.001; ns, not significant. Glc, glucose.

To test whether SIRT2 regulates β-cell proliferation, we used a pharmacological approach in isolated islets. As rodent β-cell proliferation is coupled to glucose concentration^23^, cultured mouse islets provide a means of assessing potential roles of SIRT2 in basal and adaptive β-cell proliferation, with elevated glucose mimicking increased β-cell workload. Mouse islets were incubated in the presence of increasing glucose concentrations together with the SIRT2-specific inhibitor AGK2^24^ and β-cell proliferation was quantified (Figure 1B). AGK2 treatment increased β-cell proliferation during culture in 8 mM glucose but not in 5 mM or 16.8 mM glucose (Figure 1B), indicating that SIRT2 inhibits β-cell proliferation in a glucose concentration-dependent manner. To determine whether this function of SIRT2 is conserved, we measured β-cell proliferation of AGK2-treated human islets, revealing a similar pro-proliferative effect of SIRT2 inhibition at intermediate (8 mM) glucose (Figure 1B) despite glucose itself having little effect on human β-cell proliferation, as expected^25–27^. While SIRT2 protein is β-cell-enriched in human islets, it is expressed in both α-cells and β-cells in mice, suggesting it could regulate proliferation of non-β cells. Therefore, we determined the cell type specificity of SIRT2’s effect on proliferation by quantifying proliferation of insulin-negative islet cells, revealing little effect of SIRT2 inhibition on non-β cells (Figure 1C). Previous studies have suggested that artificially stimulating β-cell proliferation could cause DNA damage or impair insulin secretion^28,29^. However, AGK2 did not alter glucose stimulated insulin secretion or insulin content of mouse or human islets (Supplementary Figure 1B and C), nor did it activate the DNA damage response as indicated by γH2AX (Supplementary Figure 1D). The inhibitory effect of SIRT2 upon β-cell proliferation in human islets was reproduced by two additional SIRT2-specific inhibitors, AK-1^24^ and SirReal2^30^ (Supplementary Figure 1E). Altogether, these results demonstrate that pharmacological SIRT2 inhibition increases proliferation of mouse and human β-cells without interfering with β-cell function or affecting proliferation of other islet cell types.

SIRT2 inactivation exerts a glucose-dependent effect on β-cell proliferation, suggesting potential coupling between cellular metabolism and SIRT2 function. The requirement of SIRT2 for oxidized NAD^+^ as a cosubstrate together with the inhibitory effect of NADH^31^ upon SIRT2’s enzymatic activity provide plausible mechanistic links between glucose metabolism and SIRT2-mediated dampening of proliferation. High glucose exposure accelerates glycolysis in β-cells, which generates NADH at the expense of NAD^+^. We therefore predicted that islet culture in 16.8 mM glucose sufficiently depletes NAD^+^ to inactivate SIRT2, rendering pharmacological SIRT2 inhibition of little effect. To test this, we performed rescue experiments to replenish NAD^+^ in high glucose using supplementation with the NAD^+^ precursor, NMN, then determined the effect of SIRT2 inhibition on β-cell proliferation. In mouse islets, NMN supplementation alone decreased β-cell proliferation compared to DMSO- or AGK2-treated islets without NMN supplementation, suggesting that NAD^+^ depletion is necessary for maximum β-cell proliferation in high glucose (Figure 1D). Importantly, SIRT2 inhibition via AGK2 in the context of NMN supplementation increased β-cell proliferation over islets treated with NMN alone, suggesting that 16.8 mM glucose inactivates SIRT2 via NAD^+^ depletion (Figure 1D). Taken together, these results show that SIRT2 modulates β-cell proliferation in a NAD^+^-dependent manner in both human and mouse islets, thereby coupling cellular metabolism to control of proliferation by SIRT2.

### Sirt2 *deletion enhances β-cell proliferation under hyperglycemic conditions in vivo*

The role of SIRT2 in restraining β-cell proliferation ex vivo together with the age-dependent increase of SIRT2 abundance led us to predict that SIRT2 could inhibit proliferation of adult β-cells. To test this, we conditionally inactivated *Sirt2* in β-cells of adult *Sirt2^fl/fl^*mice using the tamoxifen-inducible *Pdx1CreER* transgene (*Sirt2*^Δβ^ mice hereafter) then monitored glucose homeostasis (Figure 2A and Supplementary Figure 2A). Periodic glucose tolerance tests and measurements of ad libitum blood glucose levels over the course of one year revealed no evidence for altered glycemia or glucose tolerance in *Sirt2*^Δβ^ mice (Figure 2B and C and Supplementary Figure 2B and C), consistent with ex vivo data indicating that SIRT2 inactivation is of little effect in normoglycemic conditions (Figure 1B and Supplementary Figure 1B). Moreover, markers of β-cell identity Pdx1 and Nkx6.1 were unchanged in β-cells of *Sirt2*^Δβ^ mice compared to control mice (Supplementary Figure 2E and F). Quantification of β-cell proliferation and β-cell area showed no differences between *Sirt2*^Δβ^ and control mice (Figure 2D and Supplementary Figure 2D). Similar results were obtained following *Sirt2* deletion using the *MIP-CreER* transgene (Supplementary Figure 2G-K). Overall, these results indicate SIRT2 does not regulate basal β-cell proliferation and function in homeostatic conditions.

**Figure 2.**
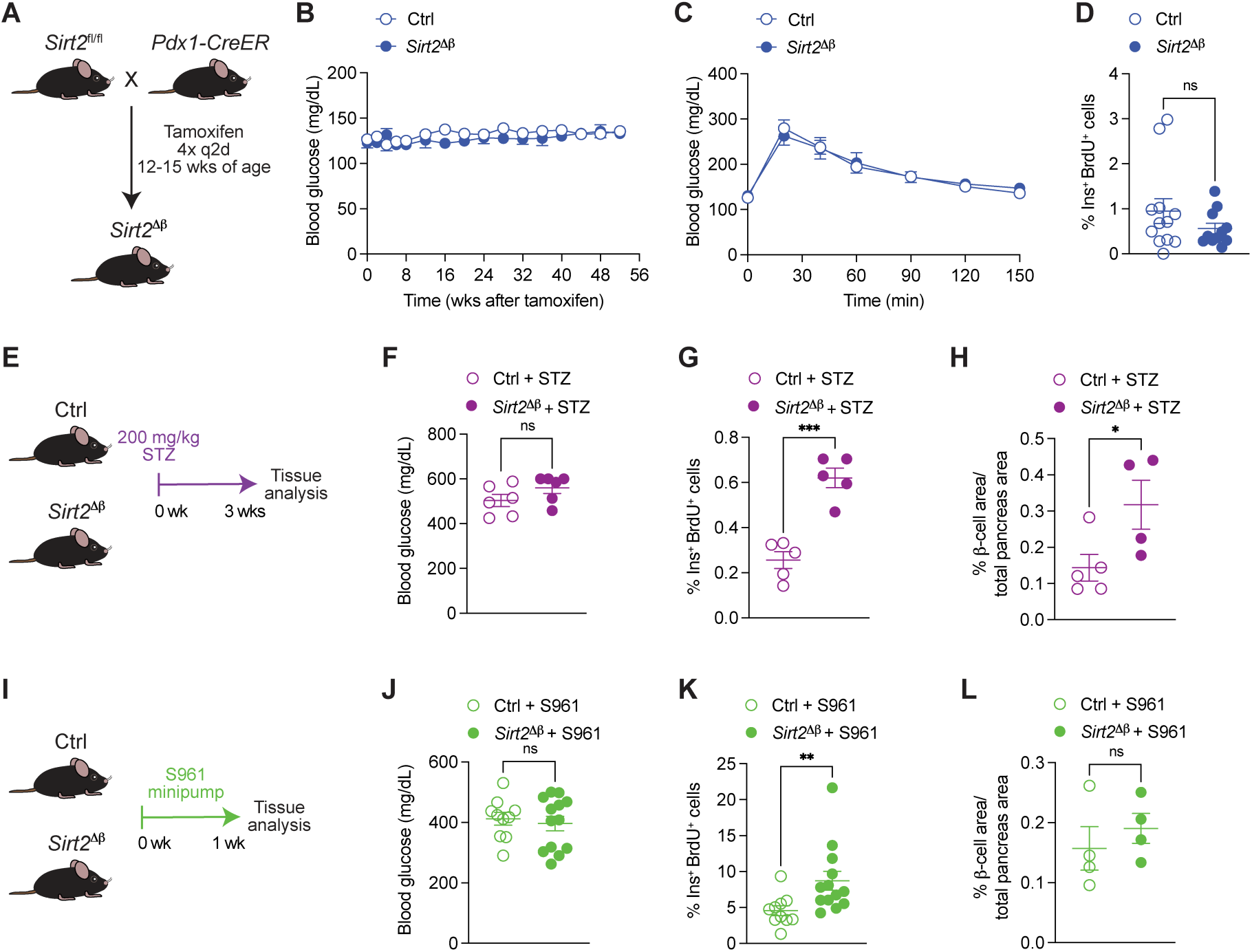
*Sirt2* inactivation enhances β-cell proliferation in hyperglycemic conditions in vivo. (**A**) β-cell-selective *Sirt2* knockout mice (hereafter called *Sirt2*^Δβ^) were generated by tamoxifen injection of 12-15-week-old *Sirt2*^fl/fl^; *Pdx1CreER* mice. *Sirt2*^+/+^ and *Sirt2*^fl/+^ mice were used as controls. (**B**) Blood glucose levels measured for 52 weeks post-tamoxifen treatment (*n* = 5-9 mice/group). (**C**) Blood glucose levels at indicated time points after an intraperitoneal glucose injection 52 weeks post-tamoxifen treatment (*n* = 5-9 mice/group). (**D**) Quantification of β-cell proliferation as percentage of insulin-positive and BrdU-positive cells relative to total insulin-positive cells 4-6 weeks following tamoxifen treatment (*n* = 11-12 mice/group). (**E**) Hyperglycemia was induced in control and *Sirt2*^Δβ^ mice by intraperitoneal injection of STZ (200 mg/kg body weight). After 3 weeks, (**F**) blood glucose levels (*n* = 6 mice/group), (**G**) β-cell proliferation (*n* = 5 mice/group) and (**H**) relative β-cell area (*n* = 4-5 mice/group) were measured. (**I**) Hyperglycemia was induced in control and *Sirt2*^Δβ^ mice by administering S961 via transplanted minipumps (20 nmol/week). After 1 week, (**J**) blood glucose levels (*n* = 10-13 mice/group), (**K**) β-cell proliferation (*n* = 10-13 mice/group), and (**L**) β-cell area (*n* = 4 mice/group) were measured. Data are shown as mean ± SEM. Statistical differences were calculated using a two-way ANOVA with Tukey post hoc analysis (B, C) or an unpaired t-test (D, F-H, J-L). \**p*<0.05, \*\**p*<0.01, \*\*\**p*<0.001; ns, not significant. q2d, every 2 days.

As we found that SIRT2 regulates β-cell proliferation in a glucose-dependent manner ex vivo (Figure 1B), we hypothesized that the effect of *Sirt2* deletion on β-cell proliferation in vivo would be contingent on hyperglycemia. To test this, we injected mice with a single high dose of streptozotocin (STZ) (Figure 2E and F) and quantified β-cell proliferation and mass. We observed increased β-cell proliferation (Figure 2G) as well as increased β-cell mass (Figure 2H) in STZ-treated *Sirt2*^Δβ^ mice compared to STZ-treated control mice. Similar findings were obtained following Sirt2 deletion using the *MIP*-*CreER* transgene (Supplementary Figure 2L-N). As a second hyperglycemic model, we administered the insulin receptor antagonist S961 to *Sirt2*^Δβ^ and control mice using micro-osmotic pumps to increase β-cell workload without ablating β-cells (Figure 2I)^32–34^. After 1 week, the mice showed highly elevated basal blood glucose with no difference in blood glucose levels between *Sirt2*^Δβ^ and control mice (Figure 2J). S961 treatment stimulated a very high rate of β-cell proliferation that was enhanced by β-cell *Sirt2* deficiency (Figure 2K). In contrast to the STZ model, *Sirt2* deletion did not increase β-cell mass compared to S961-treated control mice (Figure 2L), possibly due to the shorter duration of hyperglycemia compared to the STZ model. Altogether, these observations indicate that SIRT2 exerts context-dependent effects on β-cell proliferation in vivo, restraining adaptive proliferation during increased β-cell workload but having a minimal effect on β-cell turnover in basal conditions.

### SIRT2 deacetylates enzymes involved in oxidative metabolism of glucose and fatty acids

The above experiments demonstrate a key role for SIRT2 in dampening adaptive β-cell proliferation. To identify potential mechanisms whereby SIRT2 regulates β-cell proliferation, we sought to identify its deacetylation targets in β-cells. While SIRT2’s targets have been assessed in some contexts^22,35^, it is unclear whether these substrates are cell type- or context-dependent. We therefore cataloged proteins that are deacetylated by SIRT2 in human islets, where SIRT2 is predominantly expressed in β-cells (Figure 1A). To this end, we quantified SIRT2-dependent lysine acetylation proteome-wide by treating human islets with the SIRT2 inhibitor AGK2, enriching islet lysates with an antibody recognizing acetylated lysine, then performing mass spectrometry-based proteomics of the enriched lysates (Figure 3A). Acetylated proteins exhibiting at least 1.5-fold greater abundance in the anti-acetyl-lysine fraction of AGK2-treated relative to DMSO-treated islet lysates (64 proteins) were considered deacetylation substrates of SIRT2 (Supplementary Table 1). SIRT2 targets did not include canonical cell cycle regulators such as cyclins, cyclin-dependent kinase inhibitors, or signal transduction components downstream of known β-cell mitogens^36^. We therefore asked whether SIRT2 coordinately regulates cellular processes that could indirectly impact proliferation. Gene ontology (GO) analysis of SIRT2 substrates revealed enrichment of proteins involved in central carbon metabolism (Figure 3B and C and Supplementary Table 1) including glycolytic enzymes (GAPDH and PKM), fatty acid β-oxidation enzymes (ACAA2 and HADH), and TCA enzymes (ACO2, FH, IDH2, MDH2, and SDHA). Several of these SIRT2 targets in islets agree with SIRT2 deacetylation targets in T lymphocytes^22^, where it was shown that *Sirt2* deletion enhances enzymatic activities of GAPDH, ACO2, and SDH to augment cell proliferation. Deacetylation and consequent hyperactivation of these enzymes was suggested to contribute to increased oxidative metabolism following SIRT2 inactivation in T cells, leading us to predict that *Sirt2*^Δβ^ islets would be more oxidative than control islets. To test this possibility, we performed respirometry of *Sirt2*^Δβ^ and control islets isolated from both unchallenged and S961-treated animals. While β-cell SIRT2 inactivation did not affect oxygen consumption in islets from untreated mice (Figure 3D), islets isolated from S961-treated *Sirt2*^Δβ^ animals consumed more oxygen compared to islets from S961-treated control animals (Figure 3E). Together, these results indicate that SIRT2 deacetylates metabolic enzymes and regulates adaptive changes to islet metabolism.

**Figure 3.**
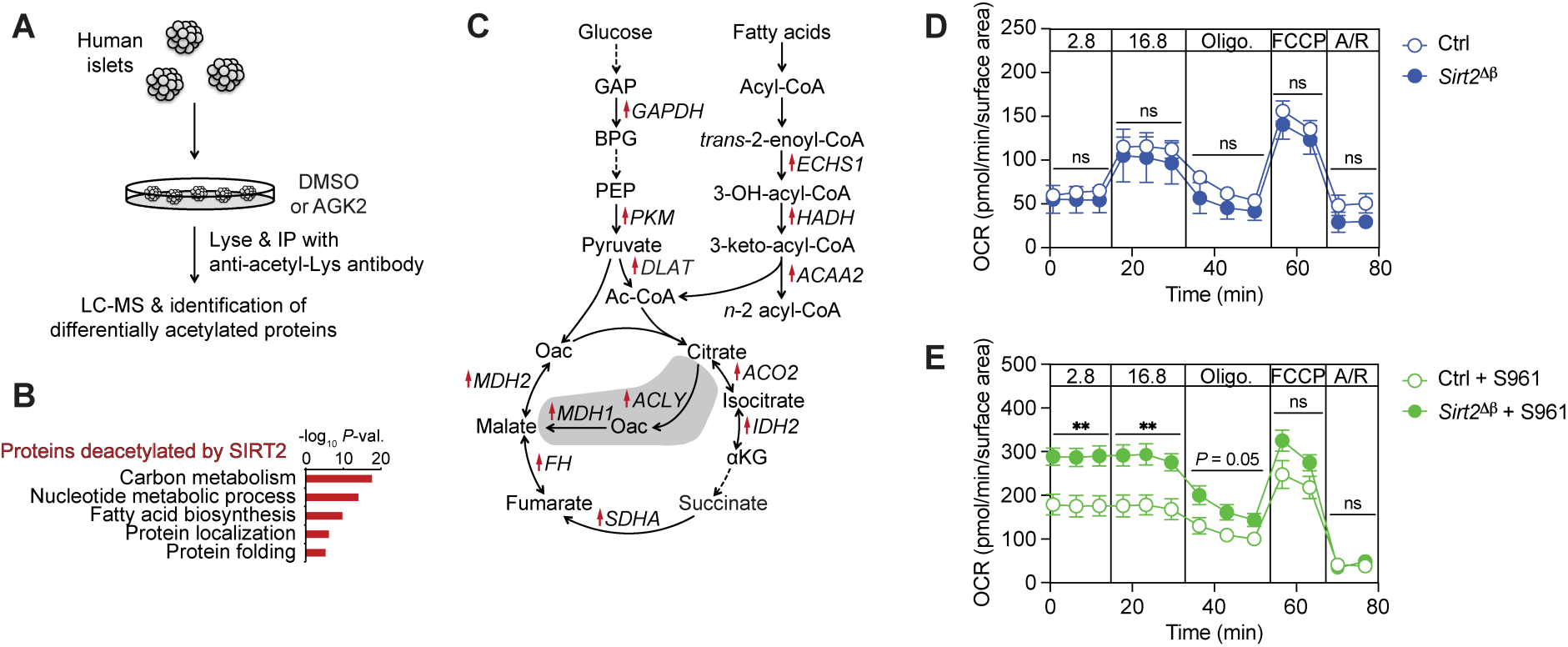
SIRT2 deacetylates metabolic enzymes and restrains islet OxPhos during hyperglycemia. **(A)** Schematic of proteomics experiment to identify proteins deacetylated by SIRT2 in human islets. (**B**) Gene ontology enrichment analysis of proteins exhibiting ≥ 1.5-fold higher acetylation following AGK2 treatment compared to vehicle control as detected by differential abundance following lysate enrichment using an antibody recognizing acetylated lysine. (**C**) Schematic of metabolic enzymes deacetylated by SIRT2. Proteins with increased lysine acetylation following AGK2 treatment are indicated with red arrows. Cytosolic reactions downstream of pyruvate are shaded. (**D** and **E**) Oxygen consumption rate (OCR) of islets from untreated (D) or S961-treated (E; as in Figure 2I) control and *Sirt2*^Δβ^ mice treated sequentially with the indicated glucose concentrations (in mM), oligomycin (Oligo.), FCCP, and antimycin/rotenone (A/R). *n* = 5-6 pools of islets/group. Data are shown as mean ± SEM. Statistical differences were calculated using a two-way ANOVA for each time block. \*\**p*<0.01 for genotype; ns, not significant. Acetyl-Lys, acetylated lysine.

### S961-induced insulin resistance and hyperglycemia remodels the β-cell transcriptional state

The glucose dependence of SIRT2’s effect upon β-cell proliferation suggests *Sirt2* deletion could affect how β-cells interpret hyperglycemia. In addition to serving as a key signal for adaptive β-cell proliferation, glucose can exert deleterious effects on β-cells through maladaptive activation of stress response pathways. To study how β-cells respond to hyperglycemic conditions in vivo, we performed single-cell RNA-seq (scRNA-seq) analysis of islets from mice treated with S961 for one week (Figure 4A). β-cells were identified based on expression of *Ins1/2* and were reclustered for further analysis, revealing five β-cell subsets with variable abundances across conditions (Figure 4B and C). We first asked how S961 impacts composition of β-cell subsets in control mice. In untreated control mice, β-cells comprised 93-95% β-1 cells and a minority population of 4-6% β-2 cells (Figure 4B and C), which agrees with the extent of transcriptional heterogeneity previously described in homeostatic conditions^37,38^. β-cell subset composition was dramatically affected by S961 treatment, which shifted β-cells to transcriptional subsets that were not detected in control islets. Islets from S961-treated mice comprised 80-88% β-5 cells and minority populations of β-3 (5-15%) and β-4 (4-5%) cells. The rapidity of these changes to β-cell subset composition far exceeds estimates of β-cell turnover during one week of S961 treatment^39^. Therefore, S961 causes β-cells to shift to a distinct set of transcriptional states that are exceedingly rare in unchallenged conditions.

**Figure 4.**
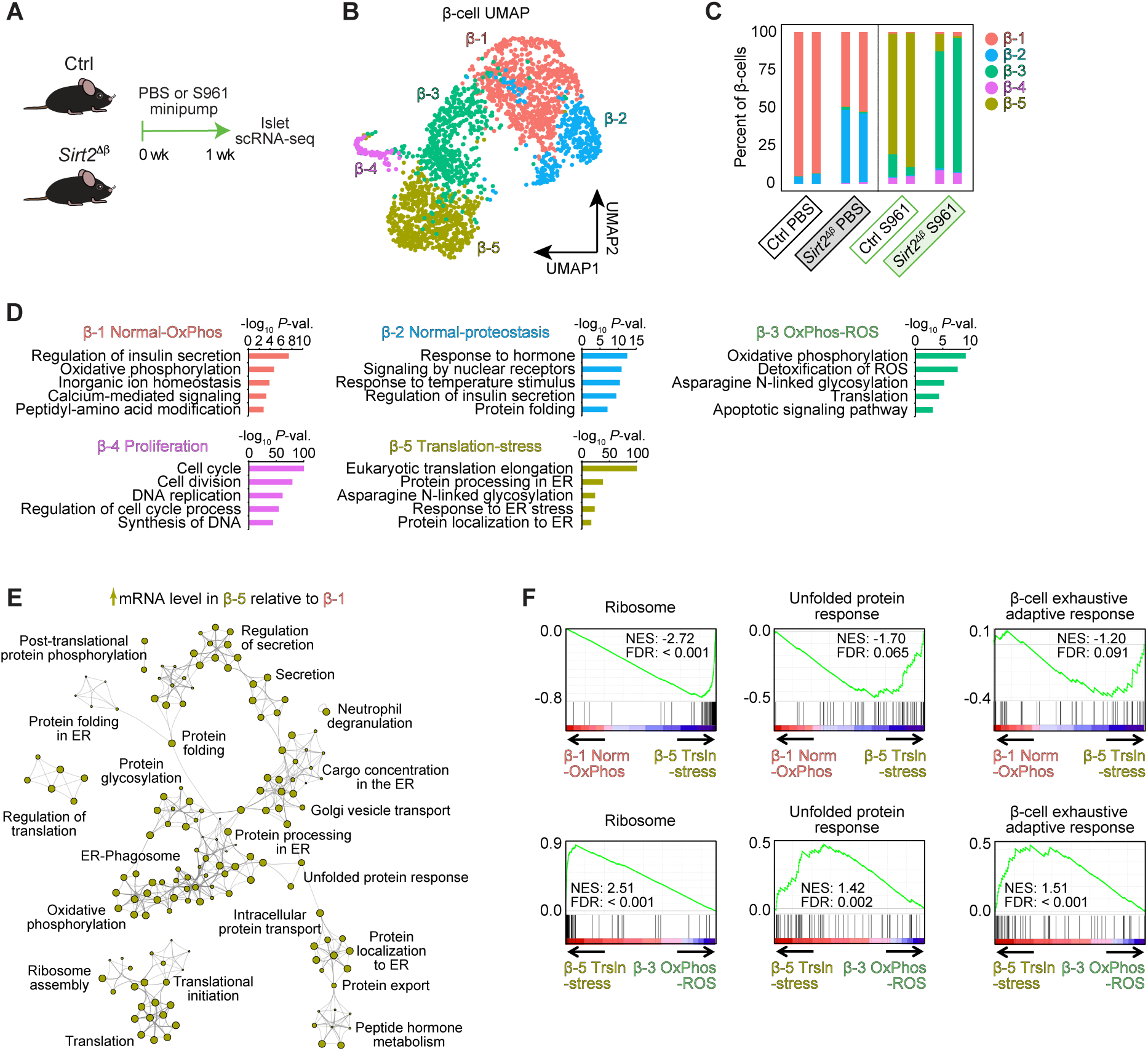
*Sirt2* inactivation affects the transcriptional response of β-cells to hyperglycemia. (**A**) Schematic of scRNA-seq experiment. (**B**) UMAP (uniform manifold approximation and projection) plot of clustered β-cells colored by subset. (**C**) Abundance of each β-cell subset as a proportion of total β-cells for the indicated treatments and genotypes (*n* = 2 technical replicates of islets pooled from 2-3 mice per group). (**D**) Enriched gene ontologies and pathways for mRNAs more highly expressed in each subset relative to all other β-cells (FDR < 0.05). (**E**) Network of gene ontologies and pathways for mRNAs more highly expressed in β-5 compared to β-1 cells (FDR < 0.05). (**F**) Gene set enrichment analysis of the indicated β-cell stress response gene sets for genes ranked by FDR for pairwise comparisons between the indicated subsets of β-cells. FDR was determined by Wilcoxon rank-sum test for subset-specific genes (D) and pairwise comparisons of β-cell subsets (E, F). OxPhos, oxidative phosphorylation; ROS, reactive oxygen species; norm, normal; trsln, translation.

We reasoned that the above-described effects of S961 on β-cell state likely reflects engagement of different cellular processes as β-cells respond to the systemic effects of hyperglycemia. To predict activation of cellular responses in each β-cell subset, we analyzed genes highly expressed in each subset relative to all other β-cells for enrichment of GOs and pathways from KEGG and Reactome databases (Figure 4D and Supplementary Table 2). As expected, the β-cell subset comprising the majority population in control mice (β-1) exhibited a gene expression signature of insulin secretion as well as enrichment of GO terms including metabolic pathways and ion channels characteristic of normal β-cells (Figure 4D). The minority population of β-cells in unchallenged mice (β-2) was also characterized by a signature of insulin secretion with additional enrichment of genes involved in protein folding (Figure 4D), which we speculate relates to the previously described signature of transiently elevated protein processing in the endoplasmic reticulum (ER)^40^. Considering relative abundances of these subsets in unchallenged mice (Figure 4C), we designated β-1 and β-2 cells as normal-OxPhos and normal-proteostasis, respectively. β-4 cells expressed genes involved in DNA replication and cell division, indicating these β-cells are proliferating as is well-documented to occur in response to increased workload^12,41^. Remaining β-cell subsets induced by S961 treatment (β-3 and β-5) both overexpressed genes associated with translation and protein glycosylation, with the majority-population β-5 cells further expressing a signature of protein processing in the ER and β-3 cells expressing a signature of OxPhos and ROS. β-3 and β-5 cells were therefore designated OxPhos-ROS and translation-stress, respectively. Given the similarity in pathway enrichments for several β-cell subsets, we directly compared transcriptomes between subsets to predict relative gene activities of pathways or cellular processes for each β-cell state. To this end, we determined enrichment of GO terms and pathways among genes differentially expressed between states. To analyze the major state transition following S961 treatment, we first compared the β-1 normal-OxPhos state to the β-5 translation-stress state (Figure 4E and Supplementary Figure 3A). Genes upregulated in the β-5 translation-stress state were involved in translation and ER-associated processes including ribosome assembly (*Rplp0*, *Rpl6*, *Rpl8*, and *Rps27a*), protein folding (*Calr* and *Hspa5*), peptide hormone metabolism (*Pcsk1*), and the unfolded protein response (*Herpud1*, *Ppp1r15a*, and *Wfs1*) (Figure 4E, Supplementary Figure 3C, and Supplementary Table 2). Genes downregulated in the β-5 translation-stress state related to β-cell function including regulation of insulin secretion, regulation of exocytosis, ion channel transport, and oxidative phosphorylation (Supplementary Figure 3A). Overall, these observations highlight increased expression of the translation machinery and engagement of ER-associated processes as β-cells adapt to hyperglycemia caused by S961.

### Sirt2 *deletion alters hyperglycemia-induced shifts in β-cell transcriptional state*

Having identified β-cell states associated with hyperglycemia, we asked whether SIRT2 inactivation affects glucose-induced β-cell state transitions by analyzing islets from *Sirt2*^Δβ^ mice treated with S961. Indeed, the abundance of the β-3 OxPhos-ROS state was enriched at the expense of the β-5 translation-stress state in S961-treated *Sirt2*^Δβ^ islets (Figure 4B-D). As β-3 and β-5 cells exhibited several overlapping GO term and pathway enrichments (i.e., translation and asparagine N-linked glycosylation), we directly compared these states using differential expression analysis between each β-cell state followed by enrichment analysis as above. Relative to β-5 cells, the β-3 OxPhos-ROS state showed lower expression of genes involved in translation and protein processing such as cytoplasmic translation, protein processing in the ER, insulin processing, and translational elongation (Supplementary Figure 3B and Supplementary Table 2). β-3 cells exhibited differential expression of OxPhos genes in both directions compared to β-5 cells, with upregulation of nuclear-encoded components and downregulation of mitochondrially-encoded components of the electron transport chain (Supplementary Figure 3B and D). Consistent with subset-specific enrichments, β-3 cells expressed higher levels of ROS detoxification genes *Txn1* and *Prdx1* compared to β-5 cells (Supplementary Figure 3E). To analyze genes involved in the cell type-specific stress response of the β-cell, we performed gene set enrichment analysis (GSEA) using recently published transcriptional signatures of the β-cell stress response^42^. Ribosomal protein genes and unfolded protein response (UPR) genes comprise major components of the recently described “β-cell exhaustive adaptive response” observed in Min6 insulinoma cells subjected to chronic treatment of the ER stress inducer cyclopiazonic acid (CPA), which inhibits Ca^2+^ transport into the ER^42^. The β-cell exhaustive adaptive response has been associated with transient growth arrest in Min6 cells^42^, providing a plausible link to altered proliferation. We therefore asked whether this response is activated in the above-described β-cell states, revealing that the S961-enriched β-5 translation-stress subset indeed displayed a signature of the β-cell exhaustive adaptive response, including upregulation of UPR and ribosomal protein genes, compared to the β-1 normal-OxPhos subset (Figure 4F). Conversely, β-3 OxPhos-ROS cells exhibited a much-reduced β-cell exhaustive adaptive response signature compared to β-5 translation-stressed cells (Figure 4F), including reductions in signatures of both the UPR (e.g., *Herpud1*, *Hspa5*, and *Wfs1*) and of ribosomal protein genes (e.g., *Rplp0*, *Rpl6*, *Rpl8*, and *Rps27a*) (Figure 4F and Supplementary Figure 3C). As *Sirt2* deletion prevented most β-cells from transitioning to the more stressed β-5 state in response to S961 treatment (Figure 4C), these observations suggest that hyperglycemia activates a stress response in β-cells in a SIRT2-dependent manner. This SIRT2-dependent stress response is reminiscent of the β-cell exhaustive adaptive response that includes signatures of the UPR and of ribosomal protein genes and has been described to coincide with ER stress-induced growth arrest in Min6 cells^42^. Notably, while SIRT2 regulates the relative abundance of transcriptionally defined β-cell states during S961 treatment, this is likely an effect not directly mediated by SIRT2, which is a protein deacetylase not involved in gene regulation. Altogether, our analysis of islet metabolism and transcriptionally-defined β-cell states builds the working model that SIRT2 deacetylates metabolic enzymes to restrain OxPhos during S961 treatment, and this direct effect of SIRT2 on metabolism is reflected by transcriptional changes indicating differences in the stress response of β-cells to hyperglycemia.

### Targeted delivery of antisense oligonucleotides against SIRT2 enhances β-cell proliferation in preclinical models of increased β-cell workload

Considering that SIRT2 inactivation promotes β-cell proliferation without interfering with insulin secretion or bypassing homeostatic set points for β-cell expansion, this enzyme is an attractive therapeutic target. However, SIRT2 is expressed in non-β cells and is an important tumor suppressor in some cell types^21^, underscoring the need for a high degree of cell type selectivity were SIRT2 to be therapeutically inactivated. As cell type-specific drug delivery remains a considerable challenge, we turned to a recently developed approach to selectively target β-cells with antisense oligonucleotides (ASO)^43^. Here, an ASO targeting a gene of interest is conjugated to the peptide hormone glucagon-like peptide-1 (GLP1). Following systemic administration, GLP1-conjugated ASO is selectively internalized by GLP1R-expressing cells, enabling cell type-specific knockdown of ASO-targeting mRNAs^43^. First, we tested the effectiveness and tissue specificity of β-cell-selective *Sirt2* knockdown by systemic administration of a GLP1-conjugated ASO targeting mouse *Sirt2* mRNA (GLP1-*Sirt2*-ASO) for three weeks. We first determined knockdown efficiency in normoglycemic mice, revealing effective 84% knockdown of *Sirt2* mRNA in islets (Figure 5A and B). Knockdown was further tested in other GLP1R-expressing tissues (kidney, heart) as well as liver, where unconjugated ASO have previously been shown to accumulate^44^. Systemic GLP1-*Sirt2*-ASO treatment had no significant effect on *Sirt2* expression in liver, kidney, or heart (Figure 5B), suggesting relative islet specificity of the GLP1-*Sirt2*-ASO. Furthermore, we observed no difference in islet *Glp1r* mRNA expression between control and GLP1-*Sirt2*-ASO-treated mice, indicating a lack of compensatory regulation of *Glp1r* expression following binding of the GLP1-conjugated ASO to GLP1R (Figure 5C). These results establish proof-of-principle for islet-selective *Sirt2* knockdown using an GLP1-conjugated ASO.

**Figure 5.**
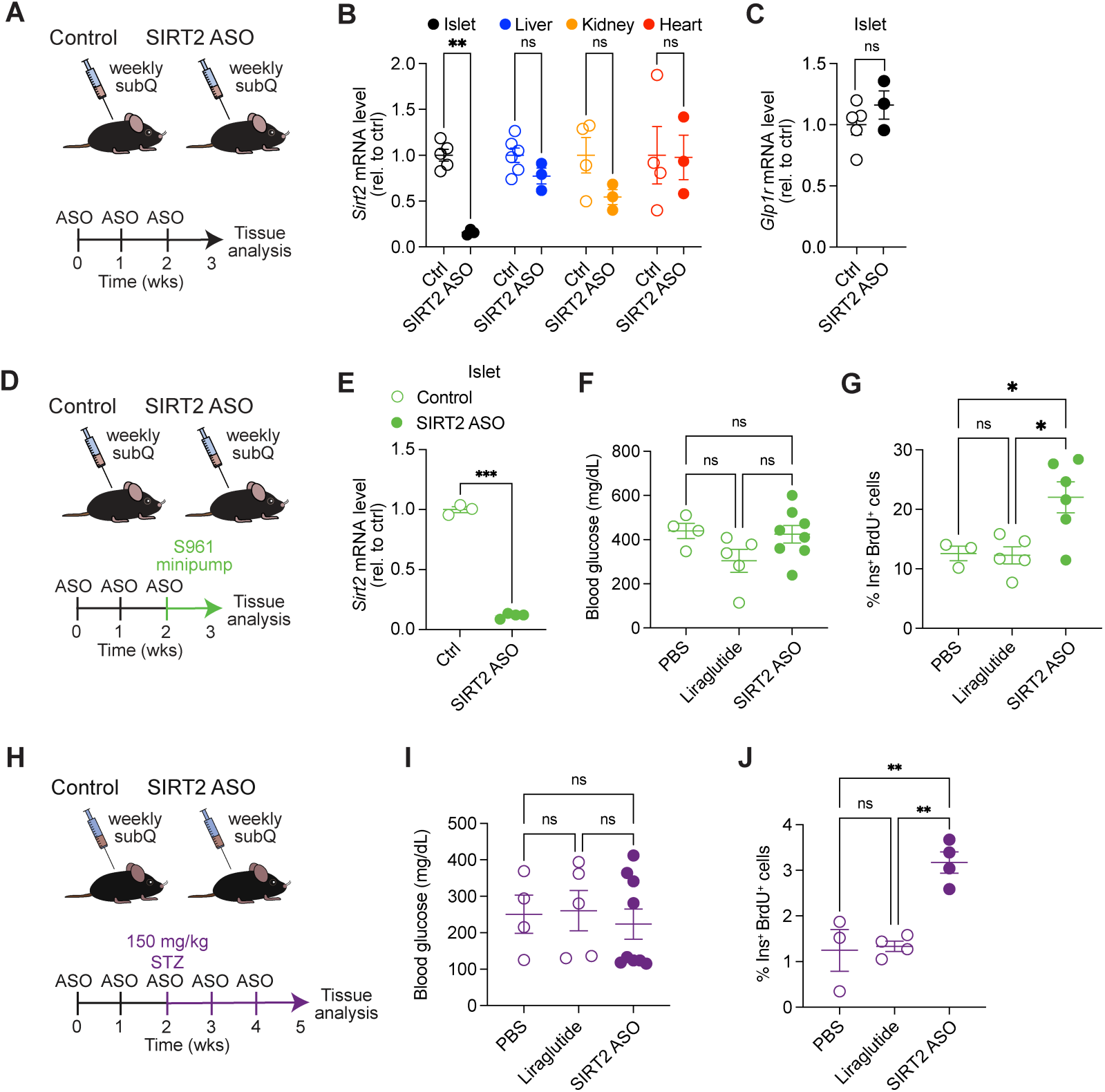
β-cell-targeted *Sirt2* ASO enhances β-cell proliferation under hyperglycemic conditions in vivo. **(A**) Schematic of systemic GLP1-*Sirt2*-ASO treatment. (**B** and **C**) qPCR analysis of *Sirt2* mRNA levels in the indicated tissues (B) and *Glp1r* mRNA level in islets (C). (**D**) Schematic of GLP1-*Sirt2*-ASO treatment followed by implantation of S961 pumps. (**E**) qPCR analysis of *Sirt2* mRNA level in islets from S961-treated mice. (**F** and **G**) Blood glucose levels (F; *n* = 4-8 mice/group) and β-cell proliferation (G; *n* = 3-6 mice/group) for the indicated groups of S961-treated mice. (**H**) Schematic of GLP1-*Sirt2*-ASO treatment in STZ-treated mice. (**I** and **J**) Blood glucose levels (I; *n* = 4-9 mice/group) and β-cell proliferation (J; *n* = 3-4 mice/group) for the indicated groups of STZ-treated mice. Data are shown as mean ± SEM. Statistical differences were calculated using two-way ANOVA followed by Fisher’s LSD test (B, E), unpaired t-test (C), or one-way ANOVA followed by Tukey’s post hoc test (F, G, I, J). \**p*<0.05, \*\**p*<0.01, \*\*\**p*<0.001; ns, not significant. Wks, weeks.

Having validated islet-specific *Sirt2* inactivation with GLP1-*Sirt2*-ASO, we asked whether the ASO enhances β-cell proliferation in preclinical models of diabetes. To confirm that metabolic stress associated with hyperglycemia does not interfere with the efficacy or selectivity of the GLP1-*Sirt2*-ASO, we analyzed tissue *Sirt2* and *Glp1r* expression in mice treated with the GLP1-*Sirt2*-ASO for three weeks and subsequently rendered acutely hyperglycemic by S961 (Figure 5D). As in normoglycemic mice, *Sirt2* knockdown was islet-selective and islet *Glp1r* expression was not affected by the GLP1-*Sirt2*-ASO (Figure 5E and Supplementary Figure 4A and B). We found no effect of the GLP1-*Sirt2*-ASO on blood glucose levels after S961 administration (Figure 5F) but observed a significant increase in β-cell proliferation in GLP1-*Sirt2*-ASO-treated mice (Figure 5G). This effect was not due to GLP1R activation as the GLP1R agonist liraglutide did not stimulate β-cell proliferation after S961 treatment (Figure 5G). Similarly, we found that the GLP1-*Sirt2*-ASO increased β-cell proliferation in mice rendered hyperglycemic by STZ (Figure 5H-J). These experiments demonstrate efficacy of β-cell-selective knockdown of *Sirt2* via ASOs in stimulating β-cell proliferation under hyperglycemic conditions. Altogether, our observations indicate that targeted SIRT2 inactivation achieves controlled, glucose-dependent β-cell proliferation.

## DISCUSSION

This work identifies SIRT2 as a regulator of adaptive β-cell proliferation whose inhibition increases β-cell numbers while preserving feedback control of β-cell mass by glucose. Pharmacological or genetic inactivation of SIRT2 selectively increases β-cell proliferation in the presence of elevated glucose levels, both ex vivo in mouse and human islets and in preclinical models of diabetes, with no effect on β-cell proliferation or mass in euglycemic conditions. We found that SIRT2 deacetylates enzymes involved in oxidative fuel metabolism, suggesting a metabolic mechanism for SIRT2’s effect on proliferation and distinguishing it from other drug targets proposed for therapeutic β-cell expansion. Altogether, these findings establish SIRT2 as a promising therapeutic target for safely increasing β-cell numbers in diabetes.

How is SIRT2 mechanistically coupled to environmental glucose concentrations in β-cells? Our studies indicate several plausible mechanisms underlying the glucose dependence of SIRT2’s function. First, we found that SIRT2 deacetylates mitochondrial metabolic enzymes and dampens respiration in high glucose in β-cells. SIRT2 deacetylation targets such as ACO2 and SDH are shared between β-cells and T cells, where it was shown that *Sirt2* deletion leads to enzyme hyperacetylation and increased oxidative metabolism^22^. As the mitogenic effect of glucose in β-cells is known to involve mitochondrial metabolism, ATP synthesis, and KATP channel closure^12^, we speculate that *Sirt2* deletion sensitizes β-cells to adaptive proliferation through increased activity of tricarboxylic acid cycle enzymes and enhanced glucose metabolism. In addition to its direct effect on metabolism, SIRT2 impacts hyperglycemia-induced changes to β-cell transcriptional state, providing a link between chronic elevations in glucose levels and SIRT2 function. A sustained elevation of glucose is known to activate β-cell stress pathways including oxidative stress^45^ as well as endoplasmic reticulum stress^46^ and the corresponding UPR^47,48^. Indeed, our scRNA-seq analysis of β-cells of hyperglycemic S961-treated mice identified SIRT2-dependent activation of various stress pathway signatures. Namely, *Sirt2* deletion ameliorated signatures of the UPR and of the β-cell exhaustive adaptive response during hyperglycemia, indicating SIRT2 impacts how β-cells interpret hyperglycemia as a stress. As stressors such as endoplasmic reticulum stress can arrest β-cell proliferation^42,46,49^, we propose that SIRT2-dependent stress responses restrain β-cell expansion during hyperglycemia, thereby linking SIRT2’s effect on β-cell proliferation to chronically elevated glucose. Further studies will be required to define the relative importance of altered metabolism and stress responses in the pro-proliferative effect of *Sirt2* inactivation, as well as the specific SIRT2 substrates mediating this effect.

The successful application of β-cell mitogenic compounds as a diabetes therapy will require achieving sufficient expansion of β-cell numbers to exert a biologically meaningful effect on glycemia concomitant with preservation of regulated insulin secretion and glycemic control^50^. Here we show that SIRT2 inactivation exerts a context-dependent effect on β-cell proliferation during hyperglycemia with little effect in euglycemic conditions, indicating that therapeutic SIRT2 targeting should not increase risk of hypoglycemia through excessive β-cell expansion and possibly formation of an insulinoma. Existing experimental drugs that show efficacy in stimulating β-cell proliferation in preclinical models include those targeting the TGFβ signaling pathway^5,51^, the tumor suppressor MEN1^9^, and the kinases DYRK1A^52^ and SIK1/2/3^53^. These compounds alter transcriptional regulation of the cell cycle machinery either directly^5,9,51,52^, or, in the case of the SIK1/2/3 inhibitor HG-9-91-01, via moderate activation of the UPR^53^. With the exception of HG-9-91-01, these agents increase β-cell proliferation in physiological glucose^5,9,51–54^, suggesting they circumvent feedback control of β-cell mass and could risk hypoglycemia if treatment is continued after the deficit in functional β-cell mass is corrected. Combination therapy with agents exhibiting synergism has been proposed as a strategy to maximize β-cell expansion^7^. Here, we found that SIRT2 regulates β-cell proliferation via a novel metabolic mechanism, opening the possibility for new combinatorial therapies utilizing distinct synergistic mechanisms. Moreover, the recovery of feedback regulation of β-cell proliferation by lowered glycemia, as observed in human islets treated with SIRT2 inhibitors, would reduce the risk of β-cell overgrowth during therapeutic expansion. Therefore, preclinical studies combinatorically applying β-cell mitogens should address whether inclusion of SIRT2-inactivating agents restores the glycemic set point for β-cell proliferation for treatments that otherwise circumvent this feedback mechanism.

Beyond avoidance of β-cell overgrowth, β-cell expansion therapy in diabetes should specifically increase proliferation of β-cells without causing hyperplasia of non-β-cells^50^. Here, we provide proof-of-principle for selective β-cell expansion using systemic administration of a *Sirt2*-targeting antisense oligonucleotide conjugated to GLP1, which is selectively taken up by β-cells. This targeted delivery approach mitigates the risk of unintended off-target effects of inactivating SIRT2 in tissues where it functions as a tumor suppressor^21^. While GLP1-conjugated antisense oligonucleotides against other mRNAs have shown efficacy in ameliorating β-cell ER stress in diet-induced obesity^55^ and β-cell dysfunction in hIAPP transgenic mice^56^, our work is the first to demonstrate increased β-cell proliferation via this approach. Considering the potential tumorigenic effect of stimulating off-target cell proliferation, it is of particular importance that mitogenic therapies exhibit cell type specificity. Given the here-described role of SIRT2 in restraining adaptive β-cell proliferation, this work opens the possibility of therapeutically targeting SIRT2 for controlled and selective β-cell expansion in diabetes.

## ACKNOWLEDGMENTS

We thank members of the Sander laboratory for helpful discussions. This work was supported by Larry L. Hillblom Foundation fellowship 2021-D-008-FEL (B.R.), NIH training grant T32GM008666-15 (J.R.B.), JDRF grant SRA-2021-1052-S-B (M.S. and M.W.), NIH grants DK068471 (M.S. and M.W.) and DK078803 (M.S.), and grants of the EPFL, ERC (ERC-2008-AdG-23118), and Swiss National Science Foundation (31003A-124713/1 and 31003A-140780) (J.A.). Human pancreatic islets were provided by the NIDDK-funded Integrated Islet Distribution Program (IIDP) (RRID:SCR _014387) at City of Hope, NIH Grant # 2UC4DK098085. This publication includes data generated at the UCSD IGM Genomics Center utilizing an Illumina NovaSeq 6000 that was purchased with funding from an NIH SIG grant (S10 OD026929) and data generated at the Biomolecular Mass Spectrometry Facility utilizing an LUMOS Orbi-Trap that was purchased with funding from an NIH SIG grant (S10 OD021724).

## AUTHOR CONTRIBUTIONS

M.W. and B.R. designed research studies, conducted experiments, acquired data, analyzed data, and wrote the manuscript. C.Z. and J.R.B. designed research studies, conducted experiments, acquired data, and analyzed data. D.P.P., F.L., M. Schlichting, A.R.H., B.L., T.P.P., and S.B. conducted experiments and acquired data. S.S., E.C.P., and H.Z. analyzed data. J.A. provided key reagents. O.S.S. designed research studies. M. Sander designed research studies, analyzed data, and wrote the manuscript.

## DECLARATION OF INTERESTS

The authors have declared that no conflict of interest exists.

## DATA AVAILABILITY

scRNA-seq data will be deposited in GEO prior to publication.

## METHODS

### Animal studies

The following mouse strains were used in this study: *C57BL/6* (Charles River), *Sirt2^flox^* ^57^, *Pdx1-CreER*^58^, and *MIP-CreER*^59^. β-cell specific *Sirt2* deletion was generated by crossing *Sirt2^flox^* mice with *Pdx1-CreER* or *MIP-CreER* mice and subcutaneous tamoxifen (Sigma) injection with 4 doses of 6 mg every other day in 12- to 15-week-old mice. Control *Sirt2^+/+^* or *Sirt2^flox/+^ CreER* mice were tamoxifen-injected in parallel. Post-tamoxifen time points are expressed relative to the final tamoxifen injection. Both male and female mice were used for all experiments. All mice were housed and bred in UCSD School of Medicine vivaria approved by the Association for Assessment and Accreditation of Laboratory Animal Care, following standards and procedures approved by the UCSD Institutional Animal Care and Use Committee. Mice were weaned at 4 weeks, maintained on a 12-hour light cycle, and fed ad libitum with standard rodent chow (PicoLab® Rodent Diet 20 5053). To render mice hyperglycemic, 12- to 15-week-old mice were treated with either STZ or S961 starting 3 weeks after the final tamoxifen injection. For the STZ experiment, mice were fasted for 4 hours before receiving an intraperitoneal injection of STZ (Sigma) dissolved in citrate buffer (pH 4.5) at doses of 150 or 200 mg/kg body weight for GLP1-*Sirt2*-ASO studies or for genetic *Sirt2* inactivation studies, respectively. Animals were given 10% sucrose water overnight. The insulin receptor antagonist S961 (Celtek Peptides) was administrated at 20 nmol/week by osmotic minipumps (Alzet 1007D) for 1 week. To target *Sirt2* in β-cells, 12-15-week-old *C57BL/6* mice were subcutaneously injected once weekly with 1 mg/kg GLP1-conjugated ASO^43,60^ against *Sirt2* (sequence with cEt chemistry wings underlined: <u>GTA</u>AGATACTGCA<u>CAA</u>) or a control (<u>GGC</u>CAATACGCCG<u>TCA</u>) for 3 weeks before hyperglycemia was induced via S961 or STZ as described above. Body weight and blood glucose levels were measured weekly using a glucometer (Bayer Contour glucometer; Bayer). For BrdU labeling, mice were given 0.8 mg/mL BrdU in drinking water for 7 days prior to harvesting pancreata.

### Islet isolation and islet culture

Mouse islet isolation was performed by collagenase digestion of the adult pancreas as previously described^19^. Briefly, mouse pancreata were perfused through the common bile duct with Collagenase P (Sigma) and islets were purified by density gradient centrifugation using Histopaque 1.077 (Sigma). Islets were allowed to recover from the isolation and cultured in RPMI 1640 medium (Cellgro) supplemented with 10% FBS, 8 mM glucose, 2 mM glutamine (Corning), 100 U/mL penicillin/streptomycin (Gibco), 1 mM sodium pyruvate (Cellgro), 10 mM HEPES (Gibco) and 0.25 μg/mL amphoterecin B at 37°C with 5% CO2 for 12 - 48 hours.

Human islets were received through the Integrated Islet distribution program (IIDP). Upon receipt, islets were stained with 0.02 µg/mL dithizone, 0.1 mM ammonium hydroxide solution for 10 minutes at 37°C, and hand-picked islets were cultured in CMRL1066 (Cellgro) supplemented with 10% FBS, 1.22 µg/mL nicotinamide, 1:1000 insulin-selenium-transferrin (Gibco), 16.7 µM zinc sulfate, 5 mM sodium pyruvate (Cellgro), 2 mM GlutaMAX (Gibco), 25 mM HEPES (Gibco), and 100 U/mL penicillin/streptomycin (Gibco). Detailed human islet donor information is listed in Supplementary Table 3.

Overnight recovered mouse or human islets were cultured with 0.1% DMSO (vehicle) or 9 µM AGK2 (Tocris), 25 µM AK-1 (EMD-Millipore), 10 µM SirReal2 (Tocris), 100 µM NMN (Sigma), or combinations of the above as indicated under various glucose concentrations (5 mM, 8 mM, 16.8 mM). To detect proliferating cells, islet culture media was supplemented with 10 µM EdU. After 48 hours, islets were fixed in 4% paraformaldehyde and stained as described below.

### Immunohistochemistry

Mouse pancreata were fixed in 4% paraformaldehyde at 4°C overnight. Human and mouse islet samples were fixed in 4% paraformaldehyde at room temperature for 30 minutes. After fixation, samples were washed three times with PBS and then incubated in 30% sucrose at 4°C overnight. Pancreata and islet samples were embedded with Optimal Cutting Temperature Compound (Tissue-Tek), frozen in a 100% ethanol/dry-ice bath, and sectioned at 10 µm using a Cryostat (Leica).

Mouse pancreata or human and mouse islet sections were then washed with PBS for 5 minutes and permeabilized and blocked with 1% normal donkey serum and 0.15% Triton X-100 (Fisher Scientific) in PBS for 1 hour. For detection of BrdU, antigen retrieval was performed by treatment with 2 M hydrochloric acid for 30 minutes followed by 0.1 M sodium borate for 5 minutes at room temperature prior to permeabilization and blocking. The following primary antibodies were used: guinea pig anti-insulin (Dako A0564, 1:1000), rat anti-BrdU (Novus Biologicals NB500-169, 1:250), rabbit-anti-Pdx1 (Abcam AB47267, 1:500), rabbit-anti-Nkx6.1 (LifeSpan BioSciences LS-C143534, 1:250), goat-anti-glucagon (Santa Cruz Biotechnology SC-7780, 1:1000), rabbit-anti-γH2AX (Cell Signaling 2577, 1:100). Dilutions were prepared in blocking solutions and sections were incubated overnight at 4°C. After washing, the sections were incubated with donkey-raised secondary antibodies conjugated to Alexa Fluor 488, Cy3 or Cy5 (Jackson ImmunoResearch) for 1 hour at room temperature. Nuclei were counterstained with DAPI (Sigma) at 0.1 µg/mL. EdU was detected using the Click-iT EdU Alexa Fluor 488 kit as specified by the manufacturer (Life Technologies).

Images were captured on a Zeiss Axio Observer Z1 microscope with an ApoTome module and processed with Zeiss AxioVision 4.8 software. Images were analyzed using HALO software (Indica Labs). Proliferation of β-cells was quantified as percentage of insulin-positive and EdU- or BrdU-positive cells relative to total cell numbers. For β-cell mass measurements, images covering an entire pancreas section were tiled using a Zeiss Axio Observer Z1 microscope with the Zeiss ApoTome module. The insulin-positive and total pancreas areas were measured using ImageJ and β-cell mass was calculated as insulin positive area relative to total pancreatic area.

### Insulin secretion measurements

Static insulin secretion assays were performed essentially as described^19^ after islets were allowed to recover overnight from isolation (mouse islets) or shipment (human islets). Islets were then pre-incubated for 1 hour at 37°C with CO2 in Krebs-Ringers-Bicarbonate-HEPES (KRBH) buffer (130 mM NaCl, 5 mM KCl, 1.2 mM CaCl2, 1.2 mM MgCl2, 1.2 mM KH2PO4, 20 mM HEPES pH 7.4, 25 mM NaHCO3, and 0.1% bovine serum albumin) containing 2.8 mM glucose. Afterwards, islets were size-matched and groups of 10 islets were transferred to a 96-well dish into KRBH solutions of 2.8 mM glucose or 16.8 mM glucose. After incubation for 1 hour as above, supernatant was collected, and islets were lysed overnight in a 2% HCl:80% ethanol solution. Secreted insulin and islet insulin content were determined using a mouse or human insulin ELISA kit (ALPCO). Secreted insulin was calculated as percentage of total insulin content.

### Metabolic studies

For glucose tolerance tests (GTT), fasted mice (6 hours) were injected intraperitoneally with a dextrose solution at a dose of 1.5 g/kg body weight. Plasma glucose levels were measured in blood samples from the tail vein at baseline, 20 min, 40 min, 60 min, 90 min, 120 min, and 150 min after glucose injection using a Bayer Contour glucometer (Bayer).

### Western blotting

Islets were lysed in RIPA buffer containing protease inhibitor cocktail (Roche) and phosphatase inhibitor (Sigma). Protein concentrations were determined by BCA assay (Thermo Fisher). Lysates (30 µg) were analyzed by SDS-PAGE on 12% Bis-Tris gels. Proteins were visualized after transfer onto a PVDF blotting membrane (GE Healthcare) and incubation with specific primary antibodies using horseradish peroxidase-conjugated secondary antibodies. Western blot primary antibodies included: rabbit-anti-Sirt2 (Santa Cruz Biotechnology SC-20966, 1:250), mouse-anti-β-tubulin (Sigma T5201, 1:1000). Western blot secondary antibodies included: donkey anti-rabbit HRP (VWR 95017, 1:1000) and sheep anti-mouse (GE Healthcare #NA931V, 1:1000). Protein bands were detected using Chemiluminescent Substrate (Thermo Fischer) on Blue Bio Films (Denville Scientific) with an SRX-101A developer (Konica Minolta). Densitometry analysis was performed using Image J.

### Acetylome analysis

Human islets from six donors (8000 islets/donor and condition) were cultured in 8 mM glucose with either the SIRT2 inhibitor AGK2 or DMSO control as described above, then islets were pelleted and stored at −80°C. Samples were pooled in 200 µl of 6 M Guanidine-HCl then subjected to 3 cycles of 10 minutes of boiling and 5 minutes of cooling at room temperature. The proteins were precipitated with addition of methanol to a final volume of 90% followed by vortex and centrifugation at 14000 rpm on a benchtop microfuge for 10 min. The soluble fraction was removed by flipping the tube onto an absorbent surface and tapping to remove any liquid. The pellet was suspended in 200 µl of 8 M urea, 100 mM Tris pH 8.0, then TCEP (10 mM) and Chloro-acetamide (40 mM) were added and the samples vortexed for 5 min. 3 volumes of 50 mM Tris pH 8.0 and 1:50 ratio of trypsin were added and samples were incubated at 37°C for 12 hours. Samples were then acidified with 0.5% TFA then desalted using Phenomenex Strata-X33 μm solid phase extraction columns (Part # 8B-S100_UBJ) as described by the manufacturer’s protocol. The peptides were dried by speed-vac then resuspended in 1 ml PBS. 50 μg of each anti-acetyl-lysine (acetyl-Lys) antibody-bead conjugate (AKL5C1, Santa Cruz Biotechnology and ICP0388, Immunechem) were added to the solution and mixed for 1 hour. After 3 washes with PBS the peptides were eluted using 1% TFA solution. 75% of the enriched fraction was analyzed by ultra-high-pressure liquid chromatography (UPLC) coupled with tandem mass spectroscopy (LC-MS/MS) using nano-spray ionization. The nano-spray ionization experiments were performed using an Orbitrap fusion Lumos hybrid mass spectrometer (Thermo) interfaced with nano-scale reversed-phase UPLC (Thermo Dionex UltiMate™ 3000 RSLC nano System) using a 25 cm, 75-μm ID glass capillary packed with 1.7-µm C18 (130) BEH^TM^ beads (Waters corporation). Peptides were eluted from the C18 column into the mass spectrometer using a linear gradient (5–80%) of acetonitrile with 0.1% formic acid at a flow rate of 375 μl/min for 180 min. Mass spectrometer parameters were as follows: an MS1 survey scan using the orbitrap detector (mass range (m/z): 400-1500 (using quadrupole isolation), 60000 resolution setting, spray voltage of 2200 V, ion transfer tube temperature of 275 C, AGC target of 400000, and maximum injection time of 50 ms) was followed by data dependent scans (top speed for most intense ions, with charge state set to only include +2-5 ions, and 5 second exclusion time, while selecting ions with minimal intensities of 50000 at in which the collision event was carried out in the high energy collision cell (HCD collision energy of 30%), and the fragment masses were analyzed in the ion trap mass analyzer (with ion trap scan rate of turbo, first mass m/z was 100, AGC target 5000 and maximum injection time of 35 ms). Protein identification was carried out using Peaks Studio 8.5 (Bioinformatics Solutions Inc). Proteins identified as being acetylated that also exhibited ≥ 1.5-fold differences in abundance between AGK2- and DMSO-treated human islet acetyl-Lys fractions (Supplementary Table 1) were considered differentially acetylated.

### Islet respirometry

Respirometry was performed using a protocol adapted from^61^. Briefly, islets were starved in Seahorse XF DMEM Assay Medium supplemented with 1% FBS, 2.8 mM glucose, and 5 mM HEPES (final pH of 7.4) at 37°C without CO2 for 1 hour. Following starvation, islets were seeded at a density of 10 size-matched islets per well into Matrigel-coated XFe96 Cell Culture Microplates containing Seahorse XF DMEM Assay Medium supplemented as above. The plate was then loaded into the Seahorse Bioscience XF96 Extracellular Flux Analyzer heated to 37°C, and oxygen consumption was measured sequentially for the following states: basal (2.8 mM glucose), glucose-stimulated (16.8 mM glucose), ATP synthase-independent (5 mM oligomycin), and non-mitochondrial (2 μM antimycin and 0.5 μM rotenone) respiration. Maximum uncoupled respiration was measured by treating with 1.5 μM carbonyl cyanide 4-(trifluoromethoxy)phenylhydrazone (FCCP) after the addition of oligomycin. Following respirometry, wells were imaged, and relative islet area was calculated using ZEN software. Oxygen consumption rates for each well were normalized to islet area relative to that of all wells. For normoglycemic mice, islets were cultured in standard islet medium (as described above) containing 8 mM glucose for 48 hours before performing islet respirometry. Respirometry experiments were performed the same day as islet isolation for islets isolated from S961-treated mice.

### Single cell RNA-seq data preprocessing, quality control and analysis

The 10x Genomics CellRanger analysis pipeline (*cellranger count* v2.1.0) was used for: (i) aligning the reads from the de-multiplexed FASTQ files to the *mm10* reference genome build (10x Genomics); (ii) filtering, and (iii) barcode and unique molecular identifier (UMI) counting, to generate gene expression matrices. The cell-calling algorithm in CellRanger automatically excludes barcodes that correspond to background noise, yielding a filtered gene expression matrix. The individual count matrices were further aggregated into a single feature-barcode matrix using the *cellranger aggr* function. All further downstream analysis, unless otherwise stated, was performed with R 4.0.4.

The aggregated feature-barcode matrix was imported and converted into a *SeuratObject* using the *CreateSeuratObject* function provided by Seurat v3 package. Data transformation and normalization was performed using SCTransform to fit a negative binomial distribution and regress out mitochondrial read percentage. Principal components (PCs) were then calculated and used downstream for unsupervised clustering and dimensionality reduction. Clustering was performed using the *FindNeighbors* with the first 30 PCs and the UMAP dimension reduction was then computed on the same 30 PCs using the *RunUMAP* function.

After initial clustering analysis, cells were filtered for mitochondrial reads < 5% and 1000 ≤ nFeature_RNA ≤ 5000. Polyhormonal cells were defined by setting hormone-specific thresholds for the four islet endocrine cell-types (Alpha - *Gcg*; Beta - *Ins1*; Delta - *Sst* and Gamma - *Ppy*). Based on this, polyhormonal doublets and endocrine-non-endocrine doublets were excluded from downstream analysis. PCs, neighbors, and UMAP dimensional reduction were then re-computed for the remaining cells. Based on the new PCs and neighbors, *k-means* clustering was performed and the resulting clusters were annotated based on hormone expression.

The clusters annotated as ‘Beta’ were subset from the full, aggregated data and re-normalized and re-computed to obtain a beta-specific *SeuratObject*. For this beta object, new subsets were computed at a resolution of 0.3 using *FindClusters* function, yielding five subsets (β-1 through β-5). The subset markers were identified using *FindAllMarkers* function with default parameters and subsequently used to identify functional categories from the Kyoto Encyclopedia of Genes and Genomes (KEGG) and Reactome categories with Metascape.

### RNA analysis

Total RNA was isolated from islets or homogenized tissues and purified using RNeasy columns and RNase-free DNase digestion according to the manufacturer’s instructions (QIAGEN). The quality and quantity of the total RNA was monitored and measured with a NanoDrop (NanoDrop Technologies, Inc. Wilmington) following the manufacturer’s instructions. For quantitative reverse transcription-PCR (RT-qPCR) analysis, cDNA was synthesized using the iScript™ cDNA Synthesis Kit (Bio-Rad) and 500 ng of isolated RNA per reaction. 20 ng of template cDNA per reaction and iQ™ SYBR^®^ Green Supermix (Bio-Rad) were used for real-time PCR with gene-specific primers (Supplementary Table 4) and *Tbp* as a housekeeping gene on a CFX96™ Real-Time PCR Detection System (Bio-Rad).

### Statistics

Statistical analyses were performed using Graphpad Prism 9 (GraphPad Software). Normality was tested via Shapiro-Wilk test and F-tests were performed to analyze equal variances. Data that passed both tests were analyzed by two-tailed Student’s t-test for two-group comparisons and one-way ANOVA for comparison of multiple groups (> 2) followed by Tukey’s post hoc testing. For data with multiple variables, e.g. glucose measurements over time, a two-way ANOVA for repeated measurements followed by Tukey’s post hoc test or Fisher Least Significant Difference post hoc testing was performed. All data are presented as mean ± SEM. *P* values less than 0.05 were considered significant.

## Notes

### Competing Interest Statement

The authors have declared no competing interest.

