## Supplementary Information for "Metabolic control of adaptive β-cell proliferation by the protein deacetylase SIRT2"

### Supplementary Figures

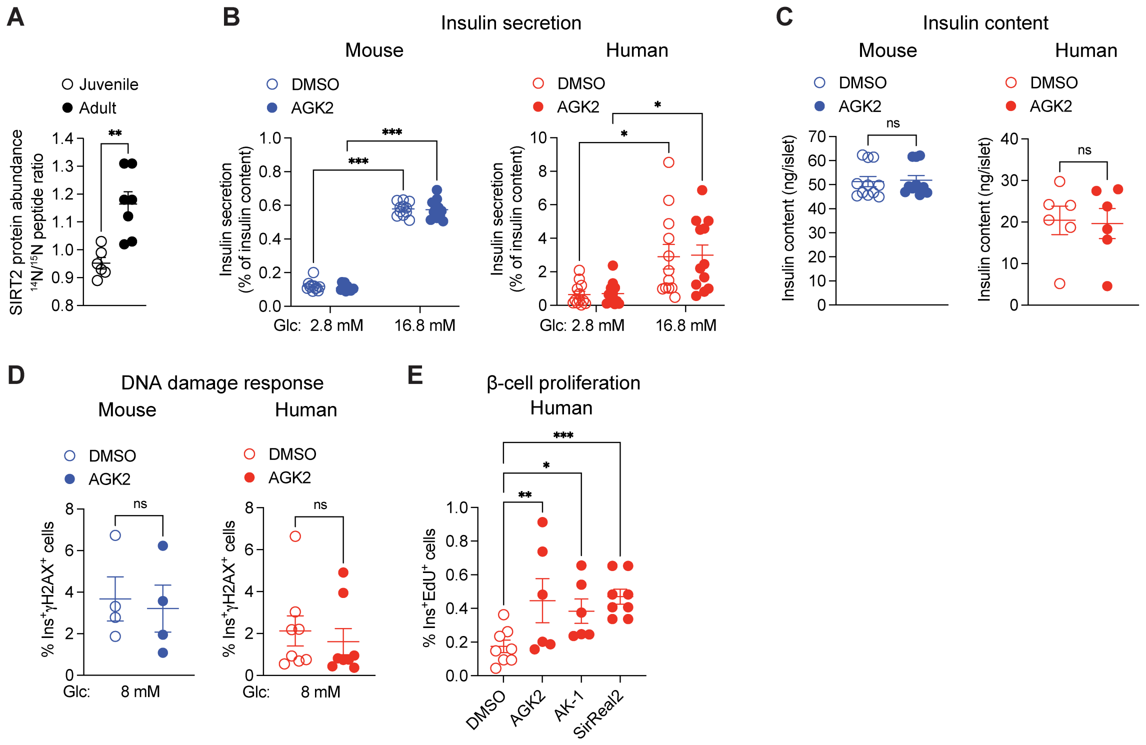


**Supplementary Figure 1. SIRT2 inhibition does not affect β-cell function.**
(**A**) Relative protein abundance for SIRT2 in islets from juvenile (one-month-old) and adult (one-year-old) mice (*n* = 4 pools of islets from distinct biological replicates/group with each peptide of SIRT2 shown as an individual data point).
(**B** and **C**) Glucose stimulated insulin secretion (B) and insulin content (C) were measured in isolated mouse (blue, *n* = 11 islet preparations/group) and human (red, B: *n* = 11, C: *n* = 6 islet preparations/group) islets after DMSO or AGK2 treatment in the indicated glucose concentrations.
(**D**) Quantification of γH2AX-positive β-cells in isolated mouse (blue, *n* = 4 islet preparations/group) and human (red, *n* = 8 islet preparations/group) islets after DMSO or AGK2 treatment in 8 mM glucose.
(**E**) Quantification of β-cell proliferation as a percentage of insulin-positive and EdU-positive cells relative to total β-cell numbers in human islets (red, *n* = 6-8 islet preparations/group) after treatment with DMSO, AGK2, AK-1, and SirReal2 in 8 mM glucose.
Data are shown as mean ± SEM. Statistical differences were calculated using unpaired t-test (A), two-way ANOVA with Tukey post hoc analysis (B), paired t-test (C, D), or one-way ANOVA with Tukey post hoc analysis (E). *$p$<0.05, **$p$<0.01, ***$p$<0.001; ns, not significant.

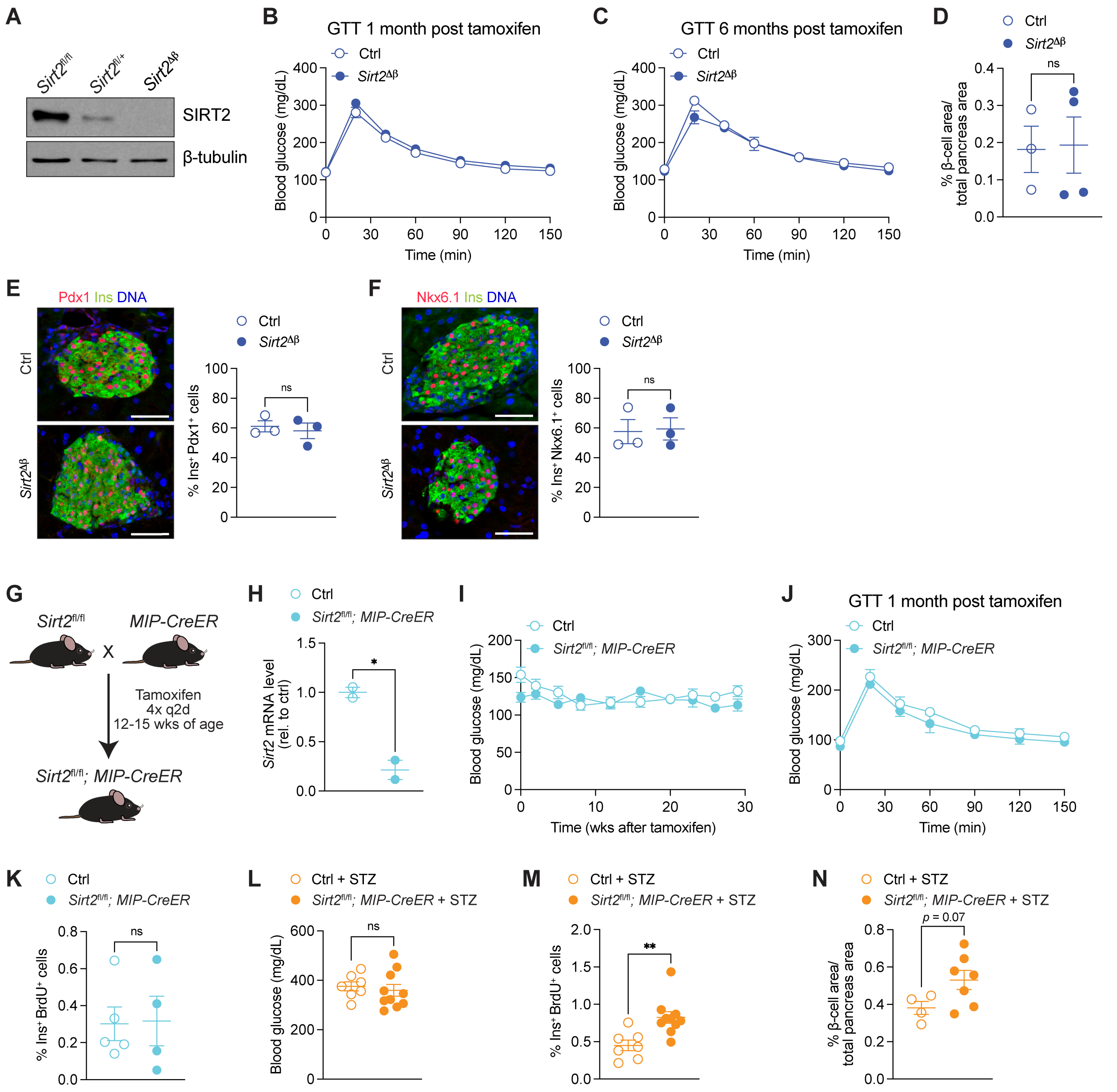


**Supplementary Figure 2. Effect of *Sirt2* deletion on glucose homeostasis and β-cell proliferation.**

(**A**) Immunoblot analysis of SIRT2 and β-tubulin in mouse islets of control or *Sirt2*$^{\Delta\beta}$ mice. Islets were pooled from 3 mice/genotype.

(**B and C**) Blood glucose levels at indicated time points after an intraperitoneal glucose injection 1 month (B) and 6 months (C) after tamoxifen treatment ($n$ = 5-9 mice/group).

(**D**) Quantification of β-cell area relative to whole pancreas area ($n$ = 3-4 mice/group).

(**E and F**) Left, representative immunofluorescence staining for the indicated proteins. Right, quantification of the percentage of β-cells expressing Pdx1 (E) or Nkx6.1 (F) for the indicated genotypes 4-6 weeks following tamoxifen treatment. DAPI was used for DNA counterstain. Scalebar is 50 μm. $n$ = 3 mice per genotype.

(**G**) Schematic for generating β-cell-specific *Sirt2* deficient mice.

(**H**) qPCR analysis of *Sirt2* mRNA level in islets from β-cell-specific *Sirt2* deficient mice relative to control mice (*n* = 2 mice/group).
(**I**) Blood glucose levels measured for 30 weeks following tamoxifen treatment (*n* = 11 mice/group).
(**J**) Blood glucose levels at indicated time points after an intraperitoneal glucose injection (*n* = 4-8 mice/group).
(**K**) Quantification of β-cell proliferation as percentage of insulin-positive and BrdU-positive cells relative to total β-cell numbers 4-6 weeks post tamoxifen treatment (*n* = 4-5 mice/group).
(**L-N**) Hyperglycemia was induced in control and β-cell-specific *Sirt2* deficient mice by intraperitoneal injection on STZ (200 mg/kg body weight). After 3 weeks, blood glucose levels (L; *n* = 7-10 mice/group), β-cell proliferation (M; *n* = 7-10 mice/group) and β-cell area (N; *n* = 4-7 mice/group) were measured.
Data are shown as mean ± SEM. Statistical differences were calculated using a two-way ANOVA with Tukey post hoc analysis (B, C, I, J) or an unpaired t-test (D-F, H, K-N). *$p<0.05$, **$p<0.01$; ns, not significant. q2d, every 2 days; wks, weeks.

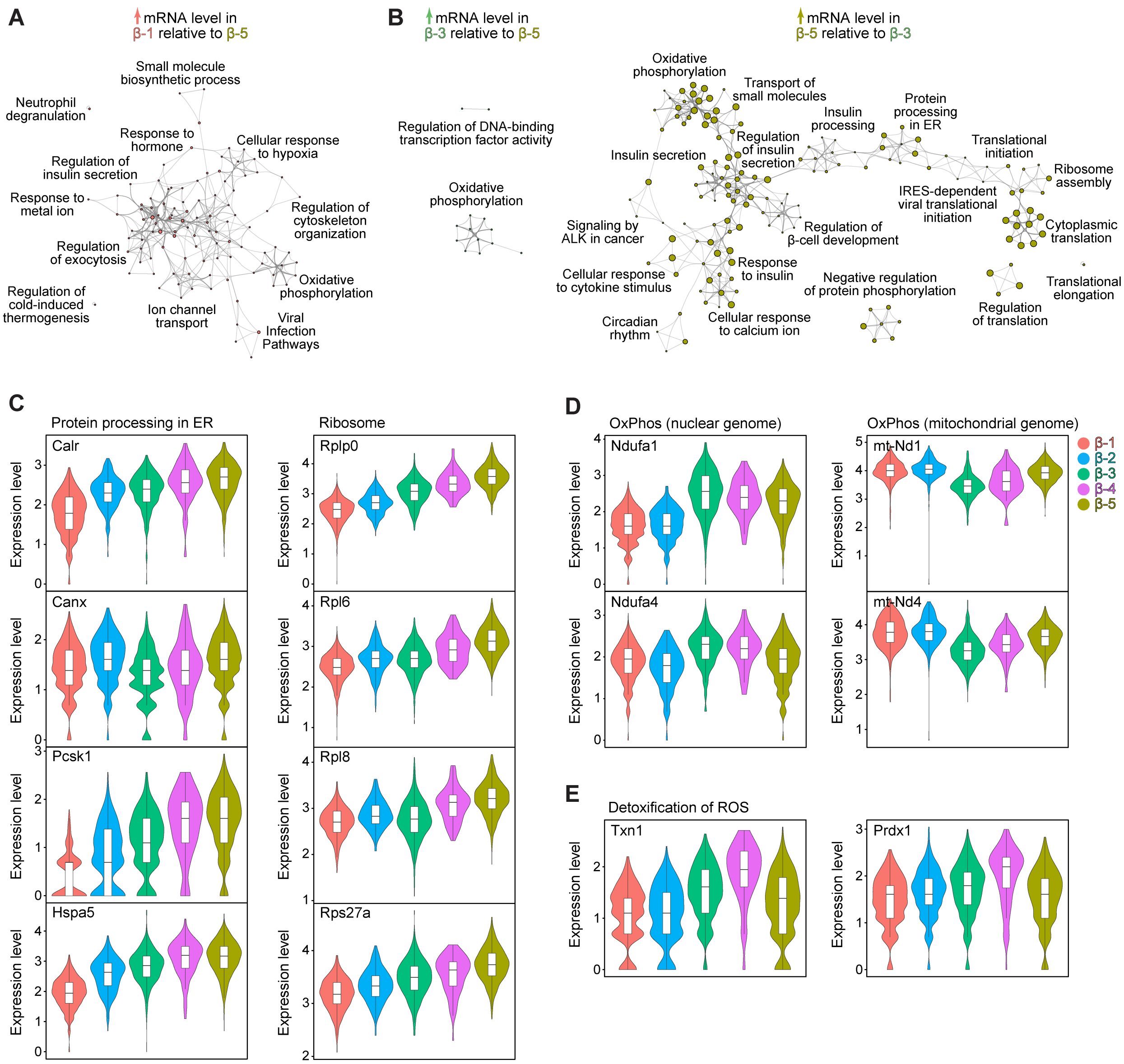


**Supplementary Figure 3. Transcriptional differences between β-cell states.**

(**A** and **B**) Networks of gene ontologies and pathways for mRNAs more highly expressed in β-1 compared to β-5 cells (A) and for mRNAs more highly expressed for the indicated comparisons (B). FDR < 0.05.

(**C-E**) Violin plots of the indicated mRNAs for each β-cell subset.

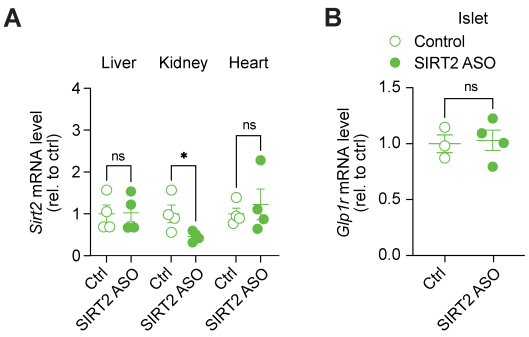


**Supplementary Figure 4. Selectivity of *Sirt2* knockdown by GLP1-*Sirt2*-ASO followed by S961 treatment.**

(**A** and **B**) qPCR analysis of *Sirt2* (A) and *Glp1r* (B) mRNA levels for the indicated tissues from S961-treated mice.

Data are shown as mean ± SEM. Statistical differences were calculated using two-way ANOVA followed by Fisher's LSD test (A) or unpaired t-test (B). *$p$<0.05, ns, not significant.

### Supplementary Tables

#### Supplementary Table 1. Acetylome analysis in human islets.

**(A)** All detected peptides and corresponding fold changes in intensity between AGK2- and DMSO-treated human islet lysates enriched for acetyl-Lys.
**(B)** Peptides identified as being acetylated from (A).
**(C)** GO term and pathway enrichment analysis of proteins whose corresponding peptide fragments were identified as being acetylated that also exhibited ≥ 1.5-fold differences in abundance between AGK2- and DMSO-treated human islet acetyl-Lys fractions.
(Supplied as Excel file: Supplementary_Table_1.xlsx)

#### Supplementary Table 2. Single cell RNA-seq analysis of control and *Sirt2*$^{\Delta\beta}$ mice treated with PBS or S961.

**(A)** mRNAs enriched in the indicated β-cell subsets relative to all other β-cells.
**(B-F)** GO term and pathway enrichment analysis of mRNAs highly expressed in each subset relative to all other β-cells (from A) for mRNAs enriched in β-1 (B), β-2 (C), β-3 (D), β-4 (E), and β-5 cells (F).
**(G and H)** Pairwise differential expression analysis of β-cell subsets comparing β-1 normal-OxPhos cells with β-5 translation-stress cells (G) or comparing β-3 OxPhos-ROS cells with β-5 translation-stress cells (H).
(Supplied as Excel file: Supplementary_Table_2.xlsx)

#### Supplementary Table 3. Human islet donor information.

Donor information, isolation center, and data generated for each human islet prep. All tissue was obtained through the Integrated Islet Distribution Program. All donors were nondiabetic.

**Human islet donor information (proliferation and GSIS experiments)**

| RRID: | SAMN08783909 | SAMN08783902 | SAMN08776506 | SAMN08776503 | SAMN08775096 | SAMN08775090 | SAMN08775023 | SAMN08775047 | SAMN08774967 | SAMN08774953 | SAMN08774820 |
|---|---|---|---|---|---|---|---|---|---|---|---|
| Age | 53 | 50 | 40 | 56 | 47 | 32 | 61 | 52 | 46 | 29 | 23 |
| Sex | M | F | M | M | F | M | F | M | F | M | F |
| BMI | 27.3 | 27.8 | 35.4 | 23.8 | 22.5 | 23.1 | 30.8 | 36.7 | 20.3 | 32.6 | 24.7 |
| HbA1c | ND | ND | ND | ND | ND | ND | ND | ND | ND | ND | ND |
| Cause of death | CV/stroke | CV/stroke | Head trauma | Head trauma | CV/stroke | Anoxia | CV/stroke | CV/stroke | CV/stroke | Head trauma | Head trauma |
| Source | Prodo | UPenn | UPenn | COH | UPenn | Prodo | Prodo | UPenn | Miami | Miami | Prodo |
| Use in study | AGK2 prolif | AGK2 prolif | AGK2 prolif | AGK2 prolif | AGK2 prolif | AGK2 prolif | AGK2 prolif | AGK2 prolif | AGK2 prolif | AGK2 prolif | AGK2, AK1/ SR2, prolif, γH2AX |

**Human islet donor information, continued (proliferation and GSIS experiments)**

| RRID: | SAMN08774480 | SAMN08774472 | SAMN08774194 | SAMN08784507 | SAMN08773854 | SAMN08930712 | SAMN08773781 | SAMN08769836 | SAMN08769806 | SAMN08769393 | SAMN08769390 |
|---|---|---|---|---|---|---|---|---|---|---|---|
| Age | 24 | 35 | 68 | 30 | 49 | 26 | 42 | 56 | 36 | 53 | 33 |
| Sex | F | M | M | F | M | M | F | F | M | F | F |
| BMI | 35.4 | 32.9 | 26.7 | 18.4 | 40.1 | 44.8 | 23.2 | 33.5 | 28.5 | 32.0 | 34.2 |
| HbA1c | ND | 5.6 | 5.3 | ND | 5.4 | 4.7 | 5.4 | 5.2 | 5.6 | 5.7 | 5.5 |
| Cause of death | CV/stroke | Head trauma | Head trauma | Anoxia | Anoxia | Head trauma | CV/stroke | Head trauma | Head trauma | CV/stroke | CV/stroke |
| Source | Miami | Miami | Prodo | UPenn | COH | Illinois | UPenn | Prodo | Miami | Prodo | Wisconsin |
| Use in study | AGK2, AK1/SR2 prolif, γH2AX | AGK2, AK1/SR2 prolif, γH2AX | AGK2, AK1/SR2 prolif, γH2AX | AGK2, AK1/SR2, NMN, non β-cell prolif, γH2AX | AGK2 prolif | AGK2, NMN, non β-cell prolif, γH2AX, GSIS | AGK2, AK1/SR2, NMN, non β-cell prolif, γH2AX, GSIS | GSIS | AGK2 prolif, GSIS | GSIS | GSIS |

### Human islet donor information, continued (proliferation and GSIS experiments)

| RRID: | SAMN08769206 | SAMN08769132 | SAMN08768969 | SAMN08769031 | SAMN08769028 | UNOS AEK1071 | SAMN08743022 | SAMN09091256 | SAMN09393858 |
|---|---|---|---|---|---|---|---|---|---|
| Age | 52 | 23 | 33 | 62 | 63 | 59 | 33 | 45 | 43 |
| Sex | F | M | M | M | F | M | F | M | F |
| BMI | 25.8 | 24.8 | 30.9 | 28.9 | 20.0 | 27.2 | 32.3 | 29.3 | 34.3 |
| HbA1c | 5.6 | 5.3 | 5.7 | 5.5 | 5.0 | 5.1 | 4.9 | 5.0 | 4.6 |
| Cause of death | CV/stroke | Anoxia | Head trauma | CV/stroke | CV/stroke | CV/stroke | Anoxia | CV/stroke | CV/stroke |
| Source | Miami | Prodo | Prodo | COH | Miami | COH | Prodo | Prodo | Prodo |
| Use in study | GSIS | AK1/ SR2, prolif, GSIS | AGK2, AGK2+/- NMN prolif | AGK2, AGK2+/- NMN prolif | AGK2, AGK2+/- NMN prolif | AGK2, AGK2+/- NMN prolif | AGK2, AGK2+/- NMN prolif | AGK2 prolif, non β-cell prolif | AGK2, AGK2+/- NMN prolif, non β-cell prolif |

ND, not determined; CV, cardiovascular; Prodo, Prodo Labs Human Islet Isolation Center; Illinois, University of Illinois; Miami, University of Miami; UPenn, University of Pennsylvania; COH, Southern California Islet Cell Resource Center at City of Hope; Wisconsin, University of Wisconsin Human Islet Core; SR2, SirReal2.

### Human islet donor information, continued (acetyl-Lys study)

| RRID: | SAMN13050553 | SAMN13049263 | SAMN13570019 | SAMN13739565 | SAMN13836615 | SAMN13972304 |
|---|---|---|---|---|---|---|
| Age | 25 | 38 | 37 | 42 | 58 | 49 |
| Sex | M | M | F | M | M | M |
| BMI | 29.3 | 24.5 | 24.0 | 37.4 | 23.3 | 34.8 |
| HbA1c | 5.3 | 5.3 | 5.2 | 5.6 | 5.7 | 5.5 |
| Cause of death | Head trauma | Anoxia | Anoxia | Anoxia | CV/stroke | CV/stroke |
| Source | UPenn | UPenn | UPenn | Prodo | Prodo | COH |
| Use in study | Acetyl-Lys | Acetyl-Lys | Acetyl-Lys | Acetyl-Lys | Acetyl-Lys | Acetyl-Lys |

ND, not determined; CV, cardiovascular; Prodo, Prodo Labs Human Islet Isolation Center; UPenn, University of Pennsylvania; COH, Southern California Islet Cell Resource Center at City of Hope.

### Supplementary Table 4. RT-qPCR primer sequences.

| mRNA | F Primer | R Primer |
|---|---|---|
| Sirt2 | GCACCTTCTACACATCACACT | ACACGATATCAGGCTTTACCAC |
| Glp1r | ACGGTGTCCCTCTCAGAGAC | ATCAAAGGTCCGGTTGCAGAA |
| Tbp | GAAGCTGCGGTACAATTCCAG | CCCCTTGTACCCTTCACCAAT |
